# Epigenetic Priming Enhances Chondrogenic Potential of Expanded Chondrocytes for Cartilage Repair

**DOI:** 10.1101/2022.10.23.513439

**Authors:** Adrienne K. Scott, Katie M. Gallagher, Stephanie E. Schneider, Abhijit Kurse, Corey P. Neu

**Affiliations:** Paul M. Rady Department of Mechanical Engineering, University of Colorado Boulder, Boulder (CO), 80309, USA; Biomedical Engineering Program, University of Colorado Boulder, Boulder (CO), 80309, USA; BioFrontiers Institute, University of Colorado Boulder, Boulder (CO), 80309, USA

**Keywords:** cartilage repair, tissue engineering, epigenetic modifications, cell fate, dedifferentiation

## Abstract

Expansion of chondrocytes presents a major obstacle in the cartilage regeneration procedure matrix-induced autologous chondrocyte implantation (MACI). Dedifferentiation of chondrocytes during the expansion process leads to the emergence of a fibrotic (chondrofibrotic) phenotype that decreases the chondrogenic potential of the implanted cells. We aim to 1) determine the extent that chromatin architecture of H3K27me3 and H3K9me3 remodels during dedifferentiation and persists when expanded chondrocytes are transferred to a 3D culture; and 2) to prevent this persistent remodeling to enhance the chondrogenic potential of expanded chondrocytes. Chromatin architecture remodeling of H3K27me3 and H3K9me3 was observed at 0, 8 and 16 population doublings in a two-dimensional (2D) culture and after encapsulation of the expanded chondrocytes in a three-dimensional (3D) hydrogel culture. Chondrocytes were treated with inhibitors of epigenetic modifiers (epigenetic priming) for 16 population doublings and then encapsulated in 3D hydrogels. Chromatin architecture of chondrocytes and gene expression were evaluated before and after encapsulation. We observed a change in chromatin architecture of epigenetic modifications H3K27me3 and H3K9me3 during chondrocyte dedifferentiation. Although inhibiting enzymes that modify H3K27me3 and H3K9me3 did not alter the dedifferentiation process in 2D culture, applying these treatments during the 2D expansion did increase the expression of select chondrogenic genes and protein deposition of type II collagen when transferred to a 3D environment. Overall, we found that epigenetic priming of expanded chondrocytes alters the cell fate when chondrocytes are later encapsulated into a 3D environment, providing a potential method to enhance the success of cartilage regeneration procedures.

## INTRODUCTION

Cartilage defects are a common result of joint trauma and sports injuries, approaching 36% of athletic populations ^1^ and 94.4% of military populations following battlefield injury ^2^. Due to the avascular nature, low cellularity, and lack of cell proliferation of chondrocytes in native cartilage, self-regeneration of the tissue is extremely limited. While proliferation of chondrocytes *in vivo* is rare, it is well established that chondrocytes proliferate when cultured *in vitro* on two-dimensional (2D) substrates. Current FDA approved cartilage tissue engineering procedures, such as matrix-induced autologous chondrocyte implantation (MACI), leverage the ability of chondrocytes to proliferate *in vitro* to repair cartilage defects. The MACI procedure involves a 2D monolayer expansion of autologous chondrocytes *in vitro* to generate sufficient cells to then transfer to a three-dimensional (3D) construct for implantation into the defect. MACI clinical trials demonstrate some efficacy when compared to other treatment methods (e.g., microfracture); however, the procedure often results in the formation of fibrocartilage ^3^. Fibrocartilage is a mechanically and biologically inferior tissue as compared to hyaline cartilage, likely due to the presence of chondrofibroblasts and related cell types which may promote formation and assembly of type I collagen-rich extracellular matrix.

While the autologous monolayer expansion process is advantageous to generate a sufficient population of chondrocytes for the MACI procedure, the process also presents a major obstacle that limits the success of the cartilage repair ^4,5^. As the chondrocytes are passaged and cultured on stiff Tissue Culture Plastic (TCP), the hyaline chondrocyte phenotype is lost and the chondrocytes dedifferentiate leading to a fibrotic phenotype ^4,6,7^, known as chondrofibroblasts. When the dedifferentiated chondrocytes are transferred to a 3D scaffold, it is well-known that the chondrocyte hyaline morphology returns ^8^. However, our previous work demonstrates that the dedifferentiated fibrotic phenotype is remembered in 3D culture and gene expression of hyaline cartilage does not return ^9^. Thus, chondrocytes are influenced by the previous 2D mechanical environment when transferred to the 3D environment, demonstrating evidence of mechanical memory ^4,10,11^. How to disrupt this memory is unknown, but if understood, this disruption could lead to methods which would increase the chondrogenic potential of expanded cells for cartilage regeneration. In this paper, we define an increase in chondrogenic potential as an increase in gene expression of chondrogenic markers (e.g., collagen II, aggrecan, or Sox9)^12^.

Increasing evidence links spatial organization of the genome to gene function and cellular memory. Recent literature, along with work from our lab, reveals that chromatin architecture maintains mechanical memories after exposure to previous non-native or stiff environments ^9,13^. Specifically, we have shown that a structural change of trimethylated H3K9 (H3K9me3) marked chromatin, an increase in H3K9me3 foci, occurs with dedifferentiation of chondrocytes and retains a memory of the dedifferentiated state, preventing redifferentiation ^9^. In addition to foci, another critical component of the nuclear architecture is the Lamina-associated domain (LAD) at the periphery of the nucleus. Genes localized in the LAD are generally repressed. The repression of non-essential genes in the LAD is critical to maintain cell-type specific functions. H3K27me3 and H3K9me3 are two histone modifications responsible for gene repression, especially in the LAD. When the enrichment of H3K27me3 and H3K9me3 in the LAD is disrupted, variable reactivation of genes can occur leading to maladaptive states, such as cancer ^14^ or cardiac pathologies ^15^. Inhibiting epigenetic modifiers presents a promising method to prevent the disruption of the LAD or general chromatin remodeling. For example, inhibiting a demethylase of H3K9me3, such as KDM4, could potentially prevent the loss of H3K9me3 toward the periphery of the nuclear envelop. Additionally, inhibiting the methyltransferase of H3K27me3, Ezh2, has previously prevented mechanically induced chromatin remodeling ^13,16^. Therefore, influencing the activity of epigenetic modifiers, such as Ezh2 and KDM4, could prevent detrimental chromatin remodeling induced by the mechanical environment from the expansion process.

The objectives of this study are to 1) explore the extent to which chondrocytes retain a memory of the dedifferentiated state encoded through the peripheral enrichment of H3K27me3 and H3K9me3, and 2) understand how disrupting the chromatin architecture changes by inhibiting the enzymes that modify H3K27me3 (treatment 1: GSK343, inhibitor of Ezh2) and H3K9me3 (treatment 2: ML324, inhibitor of KDM4 ^17^) may increase the chondrogenic potential of expanded cells for cartilage defect repair. Our data demonstrates that the peripheral enrichment of H3K27me3 and H3K9me3 decreases during chondrocyte dedifferentiation, and this reorganization is maintained when chondrocytes are transferred to a 3D hydrogel construct. We hypothesized that inhibiting epigenetic modifiers of H3K27me3 (GSK343) and H3K9me3 (ML324) would prevent chromatin remodeling during 2D culture and maintain the hyaline chondrocyte phenotype. Contrary to our hypothesis, inhibiting these epigenetic modifiers did not prevent chromatin remodeling in 2D culture. However, the epigenetic priming did alter the chondrocyte cell fate and increase the chondrogenic potential of the primed chondrocytes when transferred to a 3D hydrogel.

## MATERIALS AND METHODS

### 2D & 3D Cell Culture of Bovine Chondrocytes

Using aseptic techniques routine in our lab ^18,19^, we harvested the cartilage of the femoral medial condyles from juvenile bovine stifle joints. An institutional review board approval was not required as we did not handle live animals in this study. Briefly, we digested the tissue with 0.2% collagenase-P (Roche Pharmaceuticals, Cat. No. 11213873001) to harvest and culture primary chondrocytes. Chondrocyte media consisted of chemically defined DMEM/F12 (Gibco, Cat. No. 11330-032) supplemented with 10% Fetal Bovine Serum (Gibco, Cat. No. 26140-079), 0.1% bovine serum albumin (Sigma, Cat. No. A9418-100g), 1× penicillin streptomycin (Gibco, Cat. No. 15140-122), and 50 μg/mL ascorbate-2-phosphate (Sigma, Cat. No. 49752-10G). We seeded chondrocytes on Tissue Culture Plastic (TCP) at a density of 4×10^4^ cells/cm^2^. All samples were incubated at 5% CO_2_ and 37°C. Cells were passaged at 80% confluency. By counting cells at each passage, we determined the cell population doubled about 8 times by passage 2 and about 16 times by passage 4.

Chondrocytes were encapsulated at a concentration of 10×10^6^ cells/mL after isolation (PD0), 8 population doublings (PD8) and 16 population doublings (PD16) in 3D hyaluronan-poly(ethylene) glycol diacrylate (HA-PEGDA) hydrogels (20 mg/1mL of HA and 8.6 mg/mL PEGDA) and cultured for 1, 5, and 10 days as previously described ^9^.

### Cell Culture with Inhibitors of Epigenetic Modifiers

Twenty-four hours after seeding isolated chondrocytes (PD0), we treated the cells with media including GSK343 (1 μM, Sigma, Cat. No. SML0766-5MG), ML324 (1 μM, MedChem Express, Cat No. HY-12725), or a carrier control DMSO (0.05%) and expanded the cells to 16 population doublings. All inhibitors were dissolved in DMSO and the final concentration of DMSO in the media was maintained at 0.05%. The media of the vehicle control contained 0.05% DMSO only. After 16 population doublings, we either fixed the cells for imaging, lysed the cells in QIAzol for RT-qPCR analysis, or encapsulated the cells in HA-PEGDA hydrogels for further culture. Treated cells encapsulated in the HA-PEGDA hydrogels were then cultured in normal media to mimic the MACI procedure *in vitro* and were fixed or collected for gene expression analysis at 1, 5, and 10 days, and for analysis of protein deposition at 20 days in 3D culture.

### Live/Dead Staining

Chondrocytes cultured on TCP in the presence of epigenetic inhibitors and then encapsulated were stained and imaged to analyze cell viability. Cell viability was assessed for 2D (PD0, PD8, PD16) and 3D cultures (PD16 cells after 10 days in 3D culture) with the following applied treatments during the 2D expansion process: 1 μM ML324, 10 μM ML324, 1 μM GSK343, 2 μM GSK343 and 0.05% DMSO. Calcein AM (ThermoFisher Cat. No. C3100MP, 2 μM) and Ethidium Homodimer (ThermoFisher Cat. No E1169, 2 μM) were used to stain for live and dead cells, respectively. Cell viability was assessed qualitatively.

### Immunofluorescence Staining of 2D and 3D cultures

Cell cultures on TCP and 3D hydrogels were fixed and stained for imaging as previously described ^9^. 3D hydrogels were cut in half prior to staining to image the interior of the hydrogel. The primary antibodies used included: H3K9me3 (abcam, ab8898, 1:600) and H3K27me3 (abcam, ab6002, 1:200). Next, we incubated the cells with the corresponding secondaries including, Alexa Fluor 633 goat anti-rabbit IgG (Life Technologies, Cat. No. A21070, 1:500) and Alexa Fluor 546 goat anti-mouse IgG (Life Technologies, Cat. No. A11003, 1:500). To visualize both actin and DNA, we counterstained with 488 Phalloidin (Invitrogen, Cat. No. A12379, 1:80 in PBS) and DAPI (Invitrogen, Cat. No. D1306, 1:1000 in PBS). 3D samples cultured for 20 days were cut in half and stained for type I collagen and type II collagen. The half stained for type I collagen was fixed in 4% PFA for 30 minutes, washed, stained with the primary antibody (Abcam, Cat. No. Ab6308, 1:200) overnight and stained with the secondary for 2 hours (Abcam, Cat. No. Ab150077, 1:400). The other half was stained with the type II collagen antibody (prior to fixing with 4% PFA) by incubating the primary antibody (Invitrogen, MA5-13026, 1:100) for 2 hours followed by fixing for 30 minutes and a secondary stain for 2 hours (Abcam, Cat. No. Ab150077, 1:400). Type I and II collagen deposition was observed qualitatively through imaging.

### Confocal Imaging

We imaged all samples on an inverted Nikon A1R Confocal microscope. Specifically, we used a 60× oil immersion objective to image the nucleus/DNA (DAPI; 405nm), actin (phalloidin; 488nm), H3K27me3 (561nm) and H3K9me3 (640nm). Samples stained for type I or type II collagen were imaged with a 20× objective in the 488nm channel. All hydrogel samples were cut in half and imaged along the interior side in the hydrogel.

### Image-based Structural Nuclear Analysis and Peripheral Enrichment

Using a custom MATLAB code, we analyzed the nuclei (n > 15 nuclei/timepoint/animal) of each sample to map the location of H3K27me3 and H3K9me3 with respect to the center of the nucleus ^20^. Briefly, we used our acquired images of DAPI to determine the nuclear border and the location of the center of the nucleus. Next, we determined the normalized distances of each pixel from the center of the nucleus to bin the average marker intensities in steps of 0.1 (total of 10 bins) from the center of the nucleus (0=center, 1.0=periphery). We calculated the peripheral enrichment of both H3K27me3 and H3K9me3 by dividing the sum of the average intensity of both markers in the peripheral bins (0.7-1) by the sum of the average intensity of both markers in the center bins (0-0.3).

### Gene Expression

We isolated total RNA and reverse transcribed the RNA to cDNA from both 3D and 2D cultures to assess relative gene expression using RT-qPCR with previously described techniques ^21^. All primers were designed (sequences listed in Table 1) using NCBI Primer BLAST, ensuring that primers were separated by at least one intron and IDT performed the synthesis of primers. All data was normalized to the average of reference genes, HPRT1 and GAPDH, and relative fold changes to control samples were calculated using the ΔΔCt method.

**Table 1.**
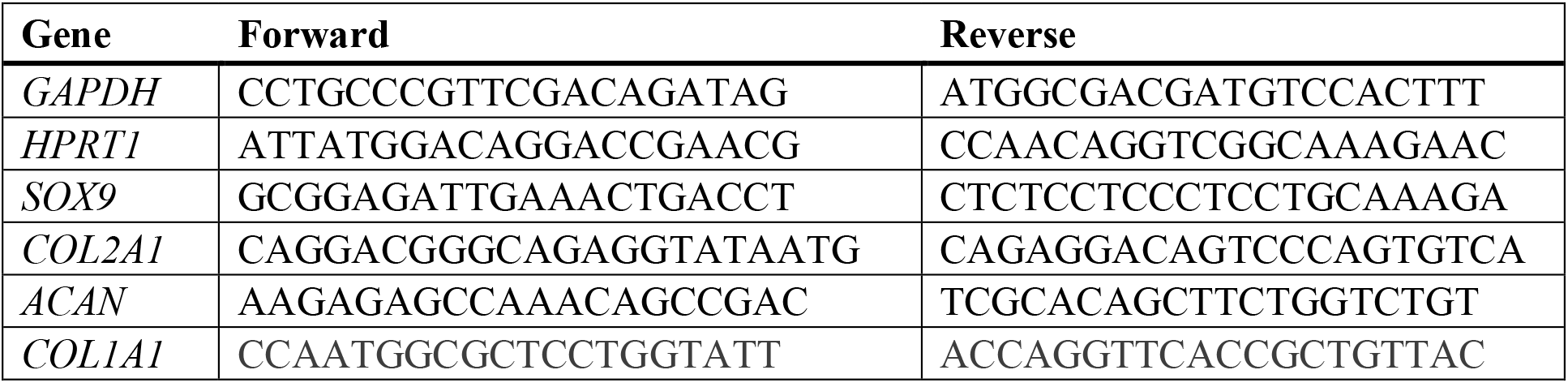
List of chondrogenic and dedifferentiation primers for gene expression analysis using RT-qPCR.

### Statistical Analysis

All datasets were analyzed using a linear mixed effect model (nlme package, version 3.1-149) with a type II or III sum of squares ANOVA (car package, version 3.0-10). Type III ANOVA was used when the model contained an interaction term, and a type II ANOVA was used for all other linear models. Passage and/or Treatment were fixed effects and animal was treated as a random effect. To meet the assumptions of the model, we evaluated the normality of the residuals using the Shapiro-Wilk test and, if necessary, the response was transformed to meet the assumptions of the model. Additionally, we evaluated the homogeneity of the residual variance for each data set by plotting the residuals against the fitted values of the linear model. In some models the variance was dependent on the passage treatment level, so a heterogenous variances model was used to allow for differences in variance among the groups. The emmeans function (package emmeans, version 1.5.3), which computes estimated marginal means, was used to test whether each treatment level was statistically significantly different from each other level. P values were adjusted for multiple comparisons using the Tukey method. P-values less than 0.05 were considered significant, and significance was denoted as *P<0.05, **P<0.01, and ***P<0.001, ****P<0.0001. We performed all statistical testing using R (version 4.0.3) and Rstudio (version 1.3.1093) software. For image data presented in Fig. 1 and Fig. 2, the experiment was repeated with 6 biological replicates and 15 to 25 nuclei were analyzed for each timepoint and replicate. For image data presented in Fig. 3 and Fig. 4, the experiment was repeated with 4 biological replicates and 18 to 25 nuclei were analyzed for each timepoint and replicate. The means reported for all gene expression experiments represent the average of 4 to 5 biological replicates. Staining of collagen type I and collagen type II at 20 days in 3D culture (Fig. 5) was completed with 4 biological replicates.

**Figure 1:**
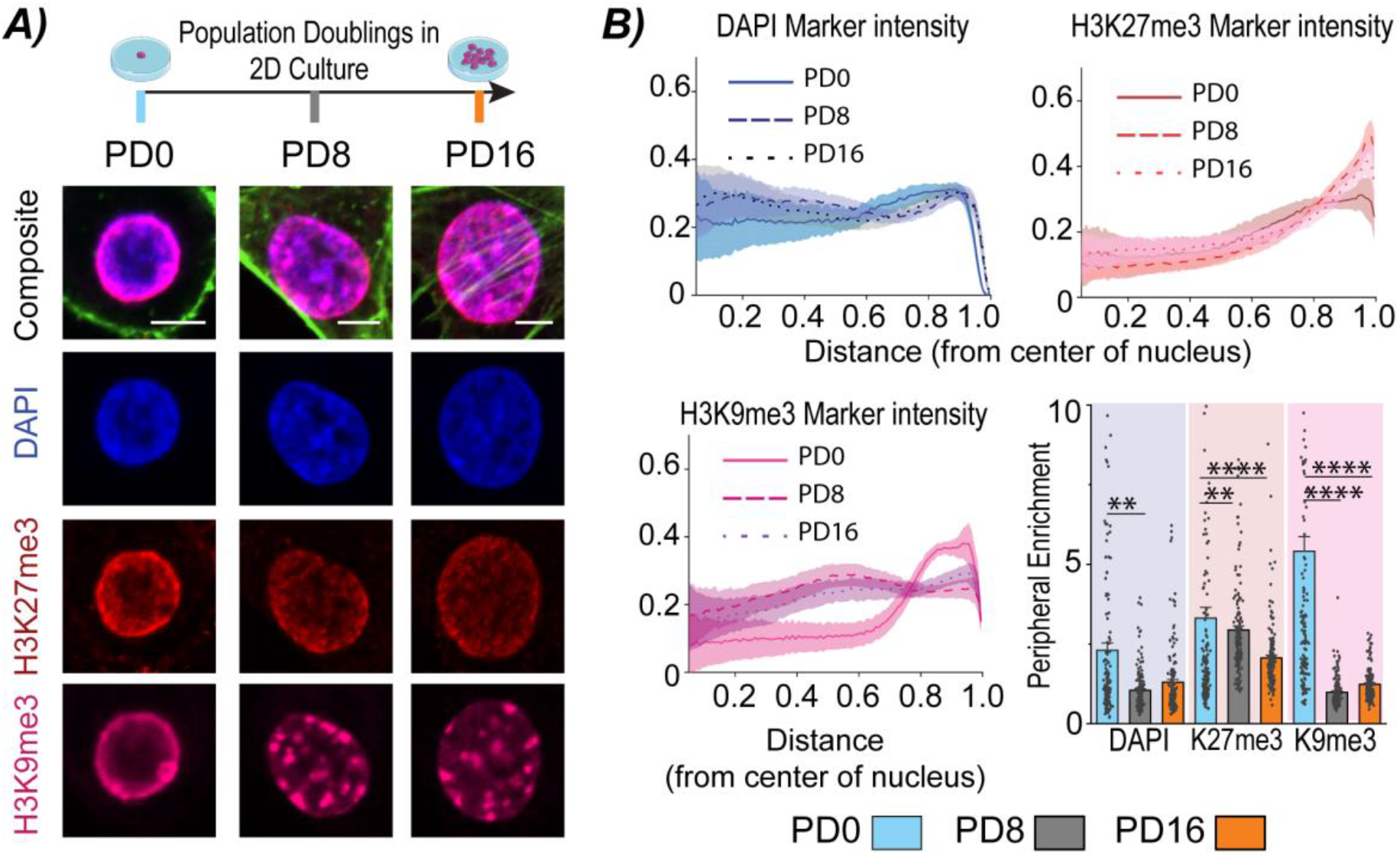
Nuclear architecture of chondrocytes, characterized by peripheral enrichment of H3K27me3 and H3K9me3, remodels during 2D cell expansion and dedifferentiation of chondrocytes. A) Chondrocytes expanded for 0 (PD0), 8 (PD8) and 16 (PD16) population doublings were counterstained for DNA (DAPI), and immunostained for H3K27me3 and H3K9me3 to visualize changes in chromatin architecture (scale= 5μm). B) Stained nuclei were analyzed to map the normalized intensity of H3K27me3 and H3K9me3 relative to the center of the nucleus (1=periphery). Peripheral enrichment was quantified by calculating the average intensity in the peripheral bin (0.7-1) divided by the center bin (0-0.3). B) Overall, peripheral enrichment of H3K27me3 and H3K9me3 decreased with increased population doublings. SEM, N=6 biological replicates, n>22 nuclei/timepoint/replicate, *p<0.05, **p<0.01, ***p<0.001, ****p<0.0001

**Figure 2:**
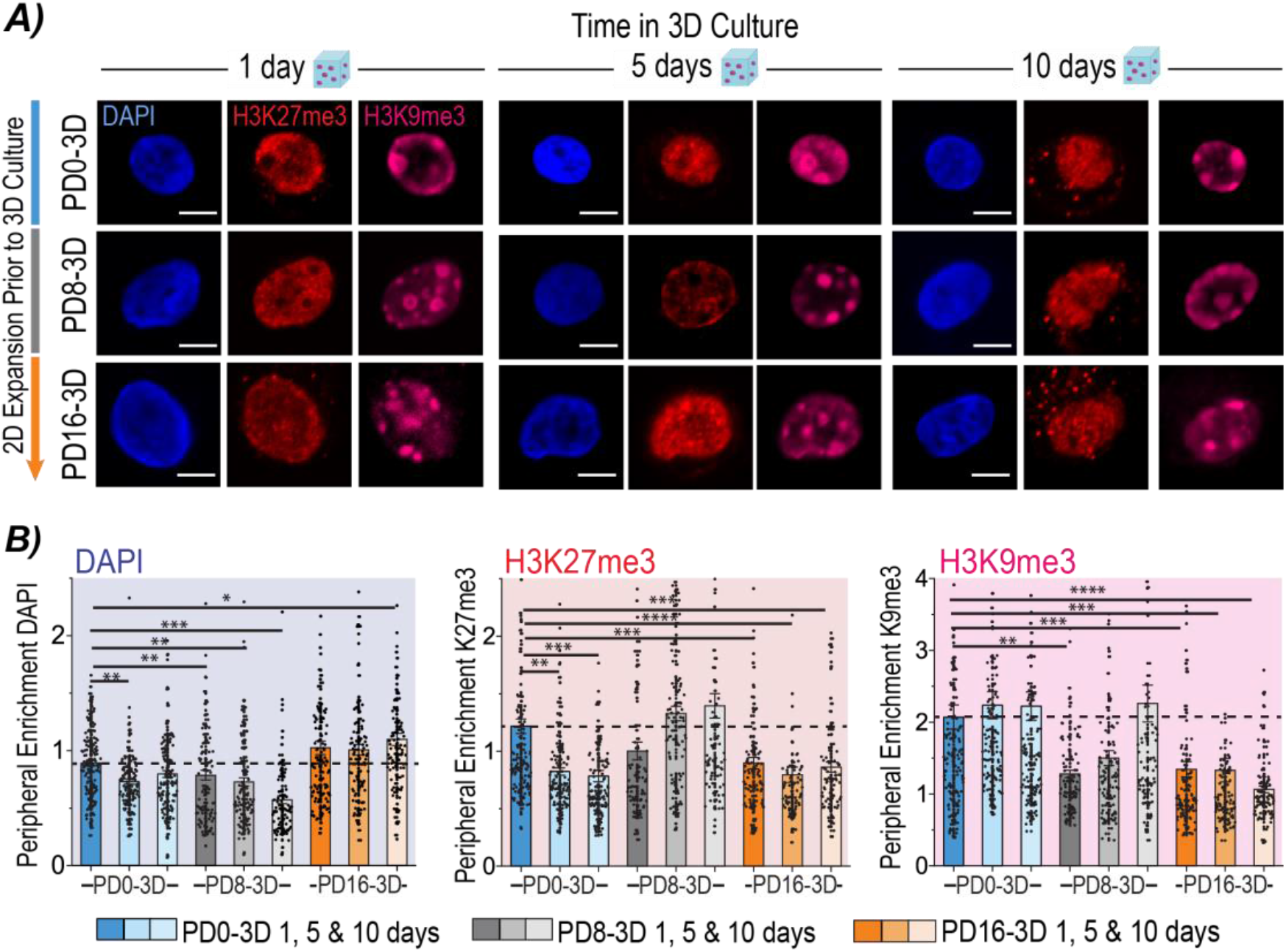
Chromatin architecture of the dedifferentiated state was restored when chondrocytes were encapsulated into 3D hydrogels, but only after a limited number of population doublings. A) Chondrocytes expanded for 0 (PD0), 8 (PD8) and 16 (PD16) population doublings were encapsulated in 3D hydrogels for 1,5, and 10 days and stained to image the chromatin architecture adaptations in 3D culture post 2D expansion. (scale= 5μm). B) Quantified peripheral enrichment of H3K27me3 and H3K9me3 was restored to levels of the PD0-3D cells only with lower population doublings (PD8-3D cells only), demonstrating a mechanical memory from the 2D expansion process. SEM, N=6 biological replicates, n>15 nuclei/timepoint/replicate, *p<0.05, **p<0.01, ***p<0.001, ****p<0.0001

**Figure 3:**
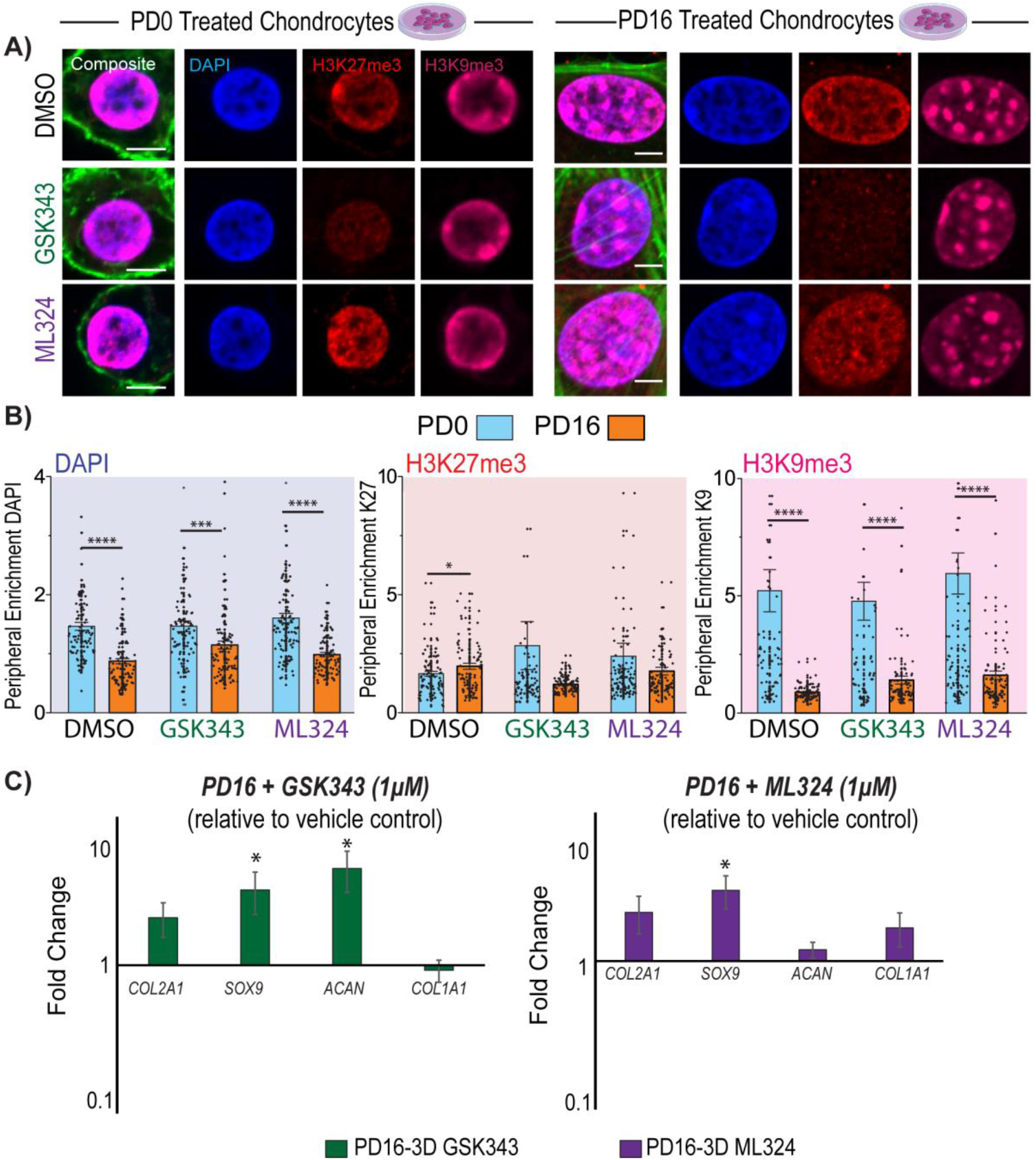
Treatments to inhibit epigenetic modifiers of H3K27me3 (GSK343) and H3K9me3 (ML324) in 2D culture did not restore the chromatin architecture of the PD0 state, but epigenetic treatments favorably increased the expression of select chondrogenic genes. A) Chondrocytes were treated with inhibitors of epigenetic modifiers of H3K27me3 and H3K9me3 during the expansion process for 16 population doublings (PD16) and imaged to assess changes in chromatin architecture (scale= 5μm). B) Epigenetic treatments did not restore peripheral enrichment levels of DAPI, H3K27me3 or H3K9me3 of PD16 cells to that of PD0 cells. SEM, N=4 biological replicates, n>18 nuclei/timepoint/replicate. C) However, select chondrogenic genes did increase for PD16 treated cells when compared to PD16 cells treated with DMSO. N=4-5 biological replicates, SEM, *p<0.05, **p<0.01, ***p<0.001, ****p<0.0001

**Figure 4:**
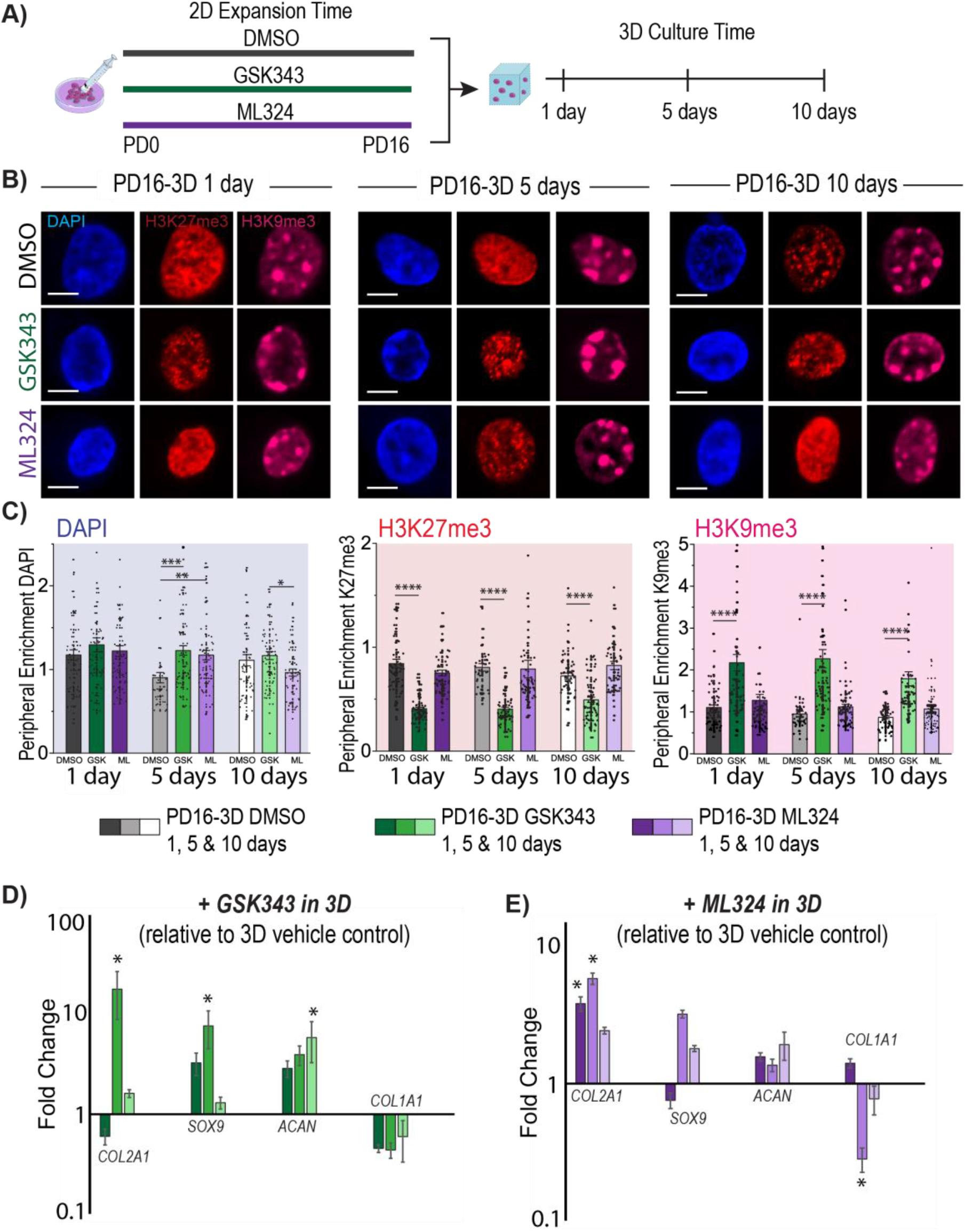
Both epigenetic treatments, GSK343 and ML324, applied during the 2D expansion process (for 16 population doublings) increased chondrogenic gene expression when encapsulated into 3D hydrogels, but only GSK343 rescued the high peripheral enrichment of H3K9me3 characteristic of native chondrocytes. A) To prevent the persistent chromatin architecture of PD16 chondrocytes, chondrocytes were primed with GSK343 or ML324 for 16 population doublings before encapsulating in hydrogels. B) Chromatin remodeling of PD16 treated cells was imaged overtime in 3D culture (PD16-3D cells). (scale= 5μm). C) While peripheral enrichment of H3K27me3 in the GSK343 treated cells remained low, the peripheral enrichment of H3K9me3 was significantly higher compared to DMSO treated PD16-3D cells for 10 days in 3D culture. Treatment with ML324 did not change peripheral enrichment levels of H3K27me3/H3K9me3 significantly when compared to DMSO controls. SEM, N=4 biological replicates, n>18 nuclei/timepoint/replicate. D) GSK343 treatment prior to encapsulation increased expression of *COL2A1*, *SOX9* and *ACAN* by 5 or 10 days in 3D culture compared to the vehicle control cells (cells treated with DMSO during 2D expansion). E) ML324 treatment prior to encapsulation increased expression of *COL2A1* and decreased expression of *COL1A1* by 5 days in culture compared to the vehicle control cells. N=4-5 biological replicates, SEM, *p<0.05, **p<0.01, ***p<0.001, ****p<0.0001

**Figure 5:**
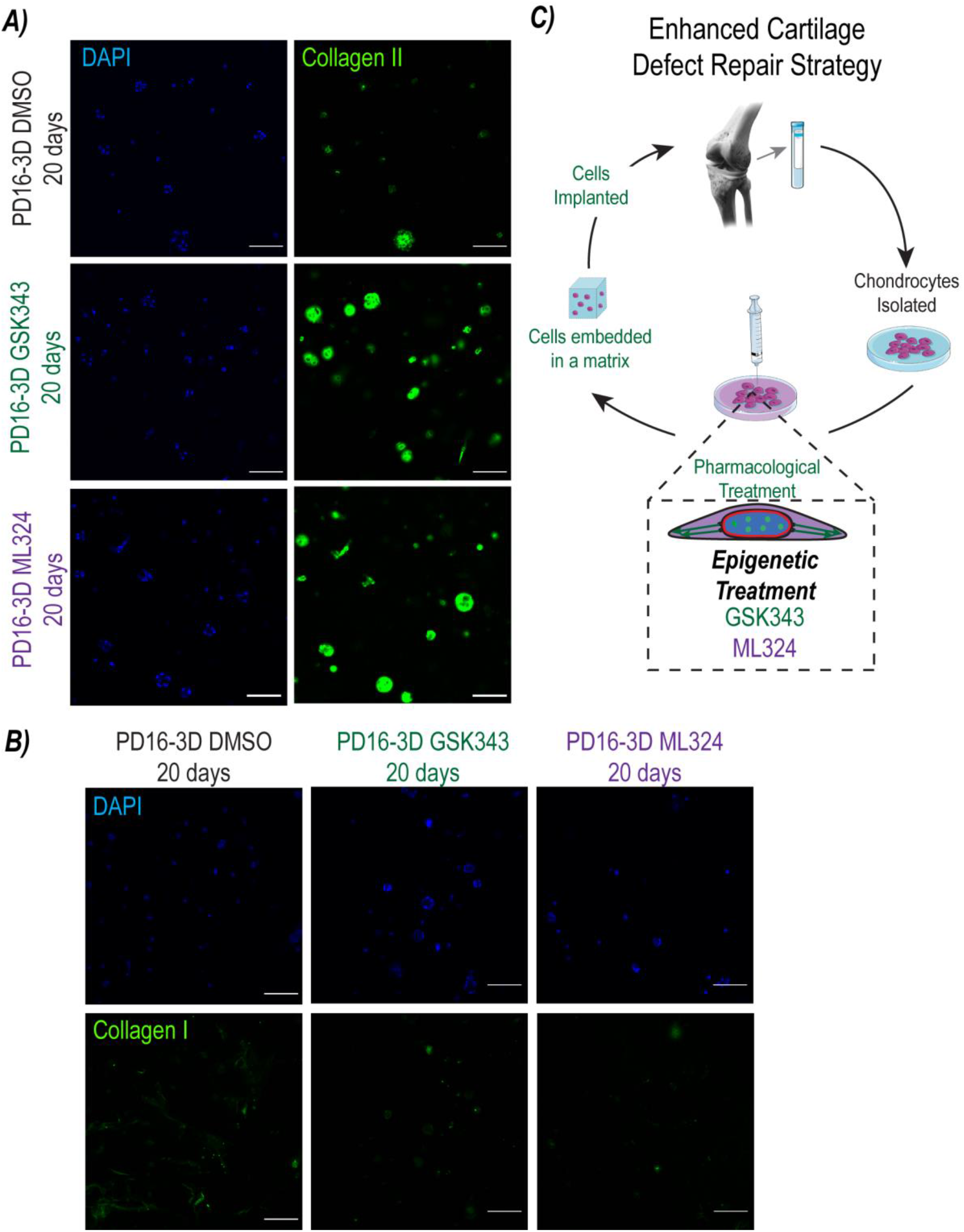
Epigenetic priming increases protein expression of type II collagen, a marker for hyaline chondrocytes, and does not increase expression of type I collagen. A) Treatment of expanded chondrocytes prior to encapsulation for 20 days increases the presence of type II collagen shown by immunostaining and imaging. N=4 biological replicates. (scale bar=100μm) B) Levels of type I collagen (shown by immunostaining and imaging), did not change for GSK343 or ML324 treated cells compared to the DMSO vehicle controls. C) We propose that the ACI/MACI procedures could be enhanced by administering a cellular treatment to chondrocytes prior to implantation into the cartilage defect to disrupt global changes in chromatin architecture and increase chondrogenic potential of treated cells.

## RESULTS

### Epigenetically marked chromatin remodels during chondrocyte dedifferentiation in 2D culture

With increased population doublings of bovine chondrocytes, the hyaline (native) chondrocyte phenotype was lost, and the chondrocytes dedifferentiated. Imaging chondrocytes during the 2D expansion process showed changes in cell morphology (cytoskeleton remodeling) characteristic of the dedifferentiation process. The native chondrocyte round morphology was lost, the area of the cell increased, and the actin cytoskeleton remodeled with increased stress fibers. These changes in cell morphology align with results from literature detailing dedifferentiation of chondrocytes ^4,22^. In addition to cell morphology changes, we also observed changes in the chromatin organization or architecture with population doublings. The structural reorganization of chromatin suggested that mechanisms altering gene regulation (e.g., histone modifications and DNA methylation) may be changing with population doublings. Specifically, we observed a shift of heterochromatin concentrated toward the periphery of the nucleus and the nucleolus to a more diffuse distribution of heterochromatin throughout the nucleus (Fig. 1A). This shift in localization of heterochromatin motivated us to explore the distribution of repressive histone modifications H3K27me3 and H3K9me3. We found that dense heterochromatin H3K27me3 and H3K9me3 regions also became more diffuse and moved towards the center of the nucleus with increased population doublings (Fig. 1A). To quantify the overall structural shift of these epigenetic changes in the nucleus, we mapped the location of these markers with respect to the center of the nucleus. We analyzed the intensity values of stained DNA (DAPI), H3K27me3 and H3K9me3 with respect to the center of the nucleus (0=center, 1=periphery) as previously described ^20^. Both H3K27me3 and H3K9me3 occupied dense regions towards the periphery of the nucleus in the PD0 cells (Fig. 1B). We calculated a peripheral enrichment score for each imaged nucleus, and found that the overall peripheral enrichment of DAPI, H3K27me3, and H3K9me3 decreased significantly with increased population doublings (Fig. 1B).

### The dedifferentiated chromatin architecture of H3K27me3 and H3K9me3 persists when chondrocytes are transferred to a 3D environment

To understand how epigenetic modifications accrued during the 2D expansion process could influence the effectiveness of cartilage defect repair procedures, we evaluated if cells retained a structural memory of the dedifferentiated state when encapsulated in a 3D culture environment. We measured changes in chromatin architecture by calculating the peripheral enrichment of DAPI, H3K27me3 and H3K9me3 and found that peripheral enrichment of DAPI and H3K27me3 decreases slightly for cells directly encapsulated into 3D hydrogels (PD0-3D: encapsulated cells without a prior expansion process), while the peripheral enrichment of H3K9me3 does not change (Fig. 2A,B). However, when cells are expanded for 8 (PD8-3D) and 16 (PD16-3D) population doublings, the low peripheral enrichment scores of H3K27me3 and H3K9me3 were retained after 1 day in 3D culture (Fig. 2A,B). The peripheral enrichment of H3K27me3 and H3K9me3 did return to the levels of PD0-3D cells by 10 days in 3D culture for cells that were only expanded to 8 population doublings (PD0-3D 1 day vs. PD8-3D 10 day, p=0.822 and p=0.9093 for peripheral enrichment of H3K27me3 and H3K9me3 respectively, Fig. 2B). In contrast, the H3K27me3 and H3K9me3 peripheral enrichment of encapsulated PD16 cells did not return to levels of the PD0-3D cells (PD0-3D 1 day vs. PD16-3D 10 day, p=0.0001 and p<0.0001 for H3K27me3 and H3K9me3 respectively) after 10 days in culture (Fig. 2B). Together, these results suggest that a memory of the remodeled chromatin structure of H3K27me3 and H3K9me3 is retained in 3D culture and how long the memory is retained depends on the previous number of population doublings in the 2D expansion.

### Inhibiting epigenetic modifiers of H3K27me3 and H3K9me3 in 2D culture does not prevent chromatin architecture remodeling, but slightly increases chondrogenic gene expression

In an effort to prevent epigenetic remodeling associated with the dedifferentiation process, we treated chondrocytes with inhibitors of epigenetic modifiers of H3K27me3 and H3K9me3, GSK343 (1μM) and ML324 (1μM) respectively, before encapsulating the chondrocytes in 3D hydrogels (Fig. 3A). Concentrations were chosen based on preliminary studies that showed more cell death when treated with high concentrations prior to 3D hydrogel encapsulation (Fig. SI 1), however we did not see a loss in cell viability in 3D culture with prior treatments of 1 μM of GSK343 and ML324 (Fig. SI 1). The chromatin architecture of PD16 cells, characterized as peripheral enrichment of H3K27me3 and H3K9me3, that were treated with GSK343 or ML324, did not differ from PD16 cells that were not treated with inhibitors. The peripheral enrichment of DAPI of the PD16 cells treated with GSK343 and ML324 all decreased compared to the PD0 cells (p=0.0007 and p<0.0001 for GSK343 and ML324 respectively, Fig. 3B), similar to the remodeling that occurred without treatment (Fig. 1B). The H3K27me3 peripheral enrichment of the PD16 treated cells did not change significantly compared to the PD0 cells, but the intensity of H3K27me3 did decrease which was expected since the methyltransferase of H3K27me3 was inhibited (Fig. 3A). The vehicle control (DMSO) did cause a slight increase in H3K27me3 enrichment of the PD16 cells compared to the PD0 cells (p=0.0278). Additionally, the peripheral enrichment of H3K9me3 of the PD16 treated cells decreased in comparison to the PD0 cells (p<0.0001 for GSK343 and ML324 treated cells). Although the epigenetic inhibitors did not significantly affect the chromatin architecture remodeling that occurs during the dedifferentiation process, the expression of select chondrogenic genes of the PD16, GSK343-treated cells did increase compared to the vehicle control cells (Fig. 3C). Treatment of PD16 cells with GSK343 led to an increase in expression of both SOX9 and ACAN compared to the DMSO treated PD16 cells (p=0.0276 and p=0.0493 for SOX9 and ACAN respectively, Fig. 3C). Additionally, the expression of SOX9 increased at PD16 population doublings for ML324 treated cells compared to vehicle control cells (p=0.0401, Fig. 3C).

### Epigenetic priming with inhibitors of H3K27me3 and H3K9me3 modifiers during 2D culture increases chondrogenic potential of expanded chondrocytes transferred to 3D culture

Although the inhibitors of the epigenetic modifiers of H3K27me3 and H3K9me3 did not alter the chromatin remodeling during the 2D culture, this epigenetic priming did alter the cell fate and chromatin architecture of the chondrocytes when the cells were transferred to a 3D hydrogel culture (Fig. 4A). Since only PD16 cells, not PD8 cells, demonstrated a persistent chromatin architecture after 3D encapsulation, we focused our analysis to PD16 cells treated with inhibitors and transferred to the 3D environment. After chondrocytes were cultured with GSK343 for 16 population doublings and transferred to a 3D culture, the peripheral enrichment of H3K27me3 remained consistently low over time as compared to DMSO controls (p<0.0001 for 1d, 5d, and 10d in culture, Fig. 4A,B,C). Interestingly, the peripheral enrichment of H3K9me3 of GSK343 primed cells increased compared to the DMSO vehicle control PD16-3D cells by day 1 in 3D culture (p<0.0001, Fig. 4B,C). This increased peripheral enrichment compared to the vehicle control cells was maintained throughout the 10 days in culture (p<0.0001 for 5d, 10ds), similar to the high H3K9me3 peripheral enrichment levels of the untreated PD0 chondrocytes (Fig. 2B). The chromatin architecture measured by peripheral enrichment of H3K27me3 and H3K9me3 did not change significantly for the ML324 treated cells compared to the DMSO vehicle control cells (all p>0.05, Fig. 4B,C).

Both treatments increased gene expression of certain chondrogenic genes. Expression of *COL2A1* and *SOX9* increased by day 5 for cells primed with the GSK343 treatment (p=0.0485 and p=0.0378 for *COL2A1* and *SOX9* respectively, Fig. 4D). By 10 days, the expression of *COL2A1* and *SOX9* for GSK343 primed cells approached levels not statistically different from the vehicle control cells, while the expression of *ACAN* remained elevated (p=0.0493, Fig. 4D). Additionally, the expression of *COL2A1* increased by day 5 in culture, while the expression of *COL1A1* decreased by day 5 in 3D culture for ML324 cells (p=0.0432 and p=0.0495 for *COL2A1* and *COL1A1* respectively, Fig. 4D). After 10 days in 3D culture, the expression of *COL2A1* and *SOX9* for the ML324 primed cells trended higher compared to controls, but the expression of *COL1A1* approached the same level as the DMSO treated control cells (Fig. 4E).

### Epigenetic priming increases protein expression of type II collagen, a marker of hyaline chondrocytes, by 20 days in 3D culture

Since gene expression of chondrogenic genes of the epigenetically primed cells generally increased compared to the DMSO control cells, we wanted to test if the corresponding protein expression of hyaline chondrocytes also increases. Because current cartilage defect repair procedures ultimately lead to a fibrocartilage repair deficient in type II collagen, we chose to analyze the presence of type II collagen in the epigenetically primed 3D samples after 20 days in 3D culture. Through observation of the images, we found that by 20 days, there was an increase in the presence of type II collagen surrounding the embedded chondrocytes in comparison to the DMSO vehicle control cells (Fig. 5A). We also assessed levels of type I collagen at 20 days in culture and found that the treatments did not change the amount of type I collagen compared to the DMSO controls (Fig. 5B). Overall, we found that epigenetic priming not only influences chondrogenic gene expression but also increases protein expression of type II collagen, a marker for the hyaline chondrocyte phenotype, presenting a possible strategy to enhance cartilage defect repair procedures like MACI (Fig. 5C).

## DISCUSSION

Regeneration of hyaline cartilage to repair cartilage defects or prevent the progression of osteoarthritis presents a major clinical challenge. Current therapies rely on a limited supply of donor tissue or lead to an inferior fibrocartilage repair ^23^. While effectiveness has been demonstrated for MACI procedures ^24^, animal model experiments suggest that the fibroblast state of the dedifferentiated chondrocyte negatively impacts the cartilage repair ^25^. We show that epigenetic priming during monolayer expansion positively alters cell fate when chondrocytes are transferred to a 3D environment and increases chondrogenic potential. Our work presents a potential solution to attenuate the long-term effects of chondrocyte dedifferentiation. Specifically, we enhanced the chondrogenic potential by treating chondrocytes during the 2D expansion process with GSK343 and ML324, inhibitors of epigenetic modifiers of H3K27me3 and H3K9me3 respectively.

Well-known work by Benya and Shaffer reported in 1982 that expanded chondrocytes dedifferentiate during monolayer culture but transferring the chondrocytes to a 3D environment (agarose gel) recovers the native chondrocyte phenotype. However, since this work was published, others reported that gene expression profiles of dedifferentiated chondrocytes are retained in 3D culture ^4,10,11^. Additionally, we have previously found that chondrogenic gene expression of cells passaged for more than 16 population doublings remains low for 10 days after transfer to a 3D culture environment. Furthermore, the work in this paper demonstrates maintenance of low peripheral enrichment of epigenetically marked chromatin after encapsulation of dedifferentiated chondrocytes. Taken together, these recent studies demonstrate that after dedifferentiation the hyaline chondrocyte phenotype does not fully return with simple encapsulation in 3D culture.

Although these studies do not fully support Benya and Shaffer’s conclusions, our data does indicate that the 3D environment is essential for increasing chondrogenic potential of dedifferentiated chondrocytes. Epigenetic treatments alone applied in 2D culture were not enough to redifferentiate the chondrocytes (Fig. 3). Epigenetic treatments in 2D culture only slightly enhanced chondrogenic gene expression. However, when treated cells were also encapsulated in 3D hydrogels the combination of epigenetic priming and the geometric constraints of the 3D environment produced an additive effect. In comparison to control cells (encapsulated cells previously treated with DMSO only), the treated cells had enhanced chondrogenic gene expression after 5 days in 3D culture and increased production of type II collagen protein after 20 days in culture. The 3D culture environment itself could influence the epigenome and chromatin architecture. For example, a previous study reports that epigenetic programing of mammary epithelial cells is influenced by 3D gel environments in comparison to 2D monolayer cultures ^26^. Additionally, recent studies using FISH have shown that relative positions of chromosomes depend on the mechanical or physical state of the nucleus ^27,28^. Since the mechanics and geometry surrounding cell environments are known to modulate nuclear morphology, it is not surprising that the native geometry (3D) would alter the chromatin state to promote cell identity.

Inhibiting the demethylase of H3K9me3, surprisingly, did not alter the peripheral enrichment of H3K9me3. We have previously found that ML324 applied at higher concentrations (10μM) to chondrocytes during the 2D expansion process causes an increase in dense regions of H3K9me3 toward the nuclear envelop and an increase in chondrogenic gene expression ^9^. Therefore, we anticipated that treating the chondrocytes with ML324 prior to encapsulation would increase the chondrogenic phenotype in 3D culture. However, in these experiments, we found that the combination of encapsulating and treating the chondrocytes with a high concentration (10μM) of ML324 led to increased cell death after encapsulation. When we decreased the concentration of ML324 to 1μM, cell viability remained high when chondrocytes were encapsulated in 3D hydrogels.

While the 1μM of ML324 did not influence the peripheral enrichment of H3K9me3, GSK343, an inhibitor of H3K27me3, did rescue the high peripheral enrichment of H3K9me3 characteristic of hyaline chondrocytes. Previous work has shown that occupation of H3K9me3 in the LAD domain decreases in response to tension while the occupation of H3K27me3 increases ^16^. Specifically, this previous study shows that force-driven enrichment of Emerin at the nuclear envelope leads to defective heterochromatin anchoring to the nuclear envelop, resulting in a switch of occupation of H3K27me3 and H3K9me3. It is possible that the opposite process is occurring due to lack of nuclear tension. We have previously shown that nuclear tension increases with cell spreading of chondrocytes during dedifferentiation in 2D culture ^29^. The restoration of H3K9me3 occupation in the LAD domain may require a decrease in levels of H3K27me3 along with a decrease in nuclear tension (i.e., transfer from a 2D to 3D environment). Alternatively, an increase in peripheral enrichment of H3K9me3 could be a compensatory mechanism to repress chromatin that is normally occupied by H3K27me3. Our results indicate that while GSK343 decreases peripheral enrichment of H3K27me3, H3K9me3 may be more enriched towards the periphery to compensate for the lack of H3K27me3. This finding is supported by a previous study in chondrocytes that shows increased levels of H3K9me3 in chondrocytes when levels of H3K27me3 decrease upon an Ezh2 siRNA knockdown ^30^.

Our previous studies support that a specific chromatin architecture is indicative of the chondrocyte cell identity and if the chromatin architecture is restored, the expression of chondrogenic genes increases. However, these results suggest that an increase in chondrogenic gene expression of select genes is possible without restoring the native chondrocyte chromatin architecture. Therefore, these results indicate that gene expression can be influenced by simply decreasing or increasing levels of H3K27me3 or H3K9me3 without large spatial reorganization of the chromatin. Additionally, there may be other changes in the chromatin architecture that are important for restoring the chondrocyte cell phenotype that are not captured solely with the assessment of peripheral enrichment. At the same time, the cells with the more characteristic chromatin architecture (GSK343 encapsulated cells) showed higher levels of expression of *COL2A1*, *SOX9* and *ACAN* when compared to ML324 treated cells. This trend indicates that high peripheral enrichment of H3K9me3 may not be necessary to increase chondrogenic gene expression, but may demonstrate that gene expression levels of chondrogenic genes could be more highly expressed with a chromatin architecture associated with the chondrocyte identity (e.g., high peripheral enrichment of H3K9me3).

Our work presents another potential approach to induce chondrogenesis of expanded chondrocytes, which could be used instead or in conjunction with previously reported methods. Previous work exploring redifferentiation of dedifferentiated chondrocytes mostly focuses on using growth factors to induce chondrogenesis ^31,32^. Future work is needed to compare the effectiveness of epigenetic priming to these other methods of chondrocyte redifferentiation, including a more complete analysis of matrix protein deposition of encapsulated epigenetically primed cells to assess the newly synthesized cartilage matrix and off target affects along with an evaluation of this method in an *in vivo* animal model. We additionally propose epigenetic priming may be used in conjunction with previously proposed methods for chondrocyte redifferentiation as treatments of epigenetic inhibitors have proven to be useful to decrease epigenetic barriers in cellular reprogramming ^33^. Just as we saw an additive effect with the 3D culture and the epigenetic treatments, it is possible treatments of GSK343 or ML324 may alter the cellular state to be more receptive to other growth factor treatments that induce chondrogenesis.

In conclusion, our study addresses a critical need to improve existing cartilage tissue regeneration methods. The use of epigenetic treatments may present a promising method to enhance MACI or ACI procedures to increase the chondrogenic potential of expanded autologous cells and could also be used in combination with other techniques to enhance the chondrogenic phenotype. Furthermore, the FDA has approved increasing numbers of epigenetic treatments ^34–37^, supporting the translational potential of this cartilage defect repair method. Future studies further evaluating our methods *in vitro* and *in vivo* are needed to understand if epigenetic priming is feasible and beneficial for cartilage defect repair procedures.

## ACKNOWLEDGEMENTS

We are grateful for the support for this work in part by grants NIH R01 AR063712 and NSF CMMI CAREER 1349735 (C.P.N.), and NIH T32 GM-065103 (S.E.S.).

## AUTHOR CONTRIBUTIONS

Conceptualization, A.K.S. and C.P.N.; Methodology, A.K.S, S.E.S, and C.P.N.; Formal Analysis, A.K.S.; Investigation, A.K.S, K.G. A.K. S.E.S; Statistical Analysis: A.K.S. Writing – Original Draft, A.K.S.; Writing – Review & Editing, All authors; Funding Acquisition, C.P.N.

## SI FIGURE

**Figure SI 1:**
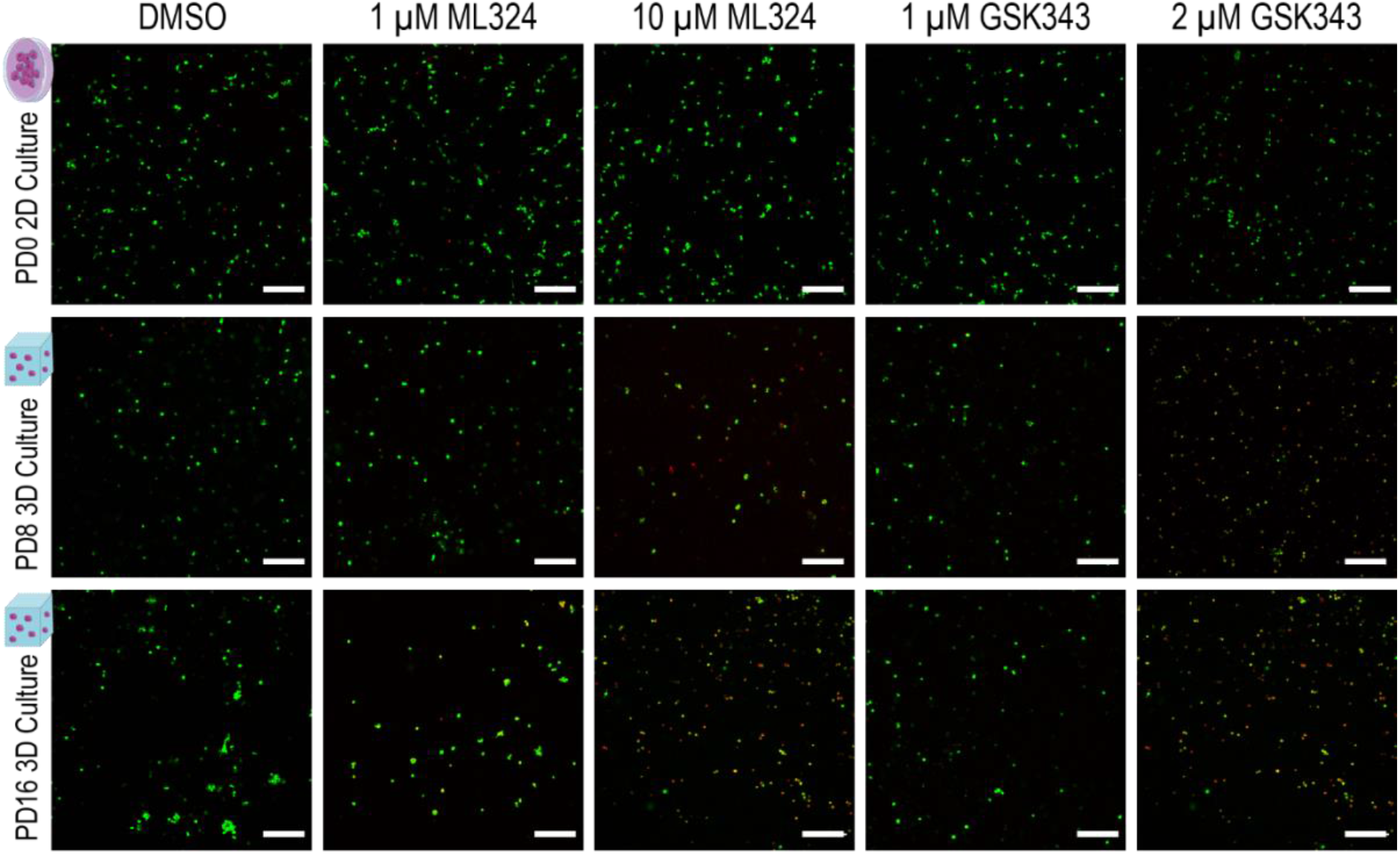
Live-dead imaging of chondrocytes treated with ML324 and GSK343 in 2D culture revealed that higher concentrations of the inhibitors influenced cell viability in 3D culture. Although we did not observe cell death with various concentrations of ML324 and GSK343 when applied in 2D culture, we observed higher cell death in 3D culture after 10 days if cells had been previously treated with 10 μM of ML324 and 2 μM of GSK343 during the 2D expansion process prior to 3D encapsulation. As a result, we treated the chondrocytes with 1 μM ML324 and 1 μM GSK343 during the expansion process to maintain high cell viability when chondrocytes were later transferred to the 3D culture. (scale bar=200μm)

## REFERENCES

1. Flanigan, D. C., Harris, J. D., Trinh, T. Q., Siston, R. A. & Brophy, R. H. Prevalence of chondral defects in athletes’ knees: a systematic review. Med. Sci. Sports Exerc. 42, 1795–1801 (2010).

2. Rivera, J. C., Wenke, J. C., Buckwalter, J. A., Ficke, J. R. & Johnson, A. E. Posttraumatic osteoarthritis caused by battlefield injuries: the primary source of disability in warriors. J. Am. Acad. Orthop. Surg. 20 Suppl 1, (2012).

3. Wegener, B. et al. Matrix-guided cartilage regeneration in chondral defects. Biotechnol. Appl. Biochem. 53, 63 (2009).

4. Ma, B. et al. Gene expression profiling of dedifferentiated human articular chondrocytes in monolayer culture. Osteoarthr. Cartil. 21, 599–603 (2013).

5. Duan, L., Liang, Y., Ma, B., Zhu, W. & Wang, D. Epigenetic regulation in chondrocyte phenotype maintenance for cell-based cartilage repair. Am. J. Transl. Res. 7, 2127 (2015).

6. Barlic, A., Drobnic, M., Malicev, E. & Kregar-Velikonja, N. Quantitative analysis of gene expression in human articular chondrocytes assigned for autologous implantation. J. Orthop. Res. 26, 847–853 (2008).

7. Schnabel, M. et al. Dedifferentiation-associated changes in morphology and gene expression in primary human articular chondrocytes in cell culture. Osteoarthr. Cartil. 10, 62–70 (2002).

8. Benya, P. D. & Shaffer, J. D. Dedifferentiated chondrocytes reexpress the differentiated collagen phenotype when cultured in agarose gels. Cell 30, 215–224 (1982).

9. Scott, A. K. et al. Epigenetic remodeling during monolayer cell expansion reduces therapeutic potential. bioRxiv 2021.12.14.472696 (2021) doi:10.1101/2021.12.14.472696.

10. Caron, M. M. J. et al. Redifferentiation of dedifferentiated human articular chondrocytes: comparison of 2D and 3D cultures. Osteoarthr. Cartil. 20, 1170–1178 (2012).

11. Kang, S.-W., Yoo, S. P. & Kim, B.-S. Effect of chondrocyte passage number on histological aspects of tissue-engineered cartilage. Biomed. Mater. Eng. 17, 269–76 (2007).

12. Schulze-Tanzil, G., Mobasheri, A., de Souza, P., John, T. & Shakibaei, M. Loss of chondrogenic potential in dedifferentiated chondrocytes correlates with deficient Shc–Erk interaction and apoptosis. Osteoarthr. Cartil. 12, 448–458 (2004).

13. Heo, S.-J. J. et al. Biophysical Regulation of Chromatin Architecture Instills a Mechanical Memory in Mesenchymal Stem Cells. Sci. Rep. 5, 16895 (2015).

14. Lochs, S. J. A., Kefalopoulou, S. & Kind, J. Lamina Associated Domains and Gene Regulation in Development and Cancer. Cells 8, (2019).

15. Joffe, B., Leonhardt, H. & Solovei, I. Differentiation and large scale spatial organization of the genome. Curr. Opin. Genet. Dev. 20, 562–9 (2010).

16. Le, H. Q. et al. Mechanical regulation of transcription controls Polycomb-mediated gene silencing during lineage commitment. Nat. Cell Biol. 18, 864–875 (2016).

17. Zhang, B. et al. KDM4 orchestrates epigenomic remodeling of senescent cells and potentiates the senescence-associated secretory phenotype. Nat. Aging 1, 454–472 (2021).

18. Novak, T. et al. In Vivo Cellular Infiltration and Remodeling in a Decellularized Ovine Osteochondral Allograft. Tissue Eng. - Part A 22, 1274–1285 (2016).

19. Neu, C. P., Khalafi, A., Komvopoulos, K., Schmid, T. M. & Reddi, A. H. Mechanotransduction of bovine articular cartilage superficial zone protein by transforming growth factor signaling. Arthritis Rheum. 56, 3706–3714 (2007).

20. Seelbinder, B. et al. Nuclear deformation guides chromatin reorganization in cardiac development and disease. Nat. Biomed. Eng. (2021) doi:10.1038/S41551-021-00823-9.

21. Barthold, J. E. et al. Recellularization and Integration of Dense Extracellular Matrix by Percolation of Tissue Microparticles. Adv. Funct. Mater. 31, 2103355 (2021).

22. Darling, E. M. & Athanasiou, K. A. Rapid phenotypic changes in passaged articular chondrocyte subpopulations. J. Orthop. Res. 23, 425–432 (2005).

23. Gudas, R. et al. A prospective randomized clinical study of mosaic osteochondral autologous transplantation versus microfracture for the treatment of osteochondral defects in the knee joint in young athletes. Arthroscopy 21, 1066–1075 (2005).

24. Rogers, B. A., David, L. A. & Briggs, T. W. R. Sequential outcome following autologous chondrocyte implantation of the knee: A six-year follow-up. Int. Orthop. 34, 959 (2010).

25. Dell’accio, F., De Bari, C. & Luyten, F. P. Molecular Markers Predictive of the Capacity of Expanded Human Articular Chondrocytes to Form Stable Cartilage In Vivo. ARTHRITIS Rheum. 44, 1608–1619 (2001).

26. Jolivet, G., Pantano, T. & Houdebine, L. M. Regulation by the extracellular matrix (ECM) of prolactin-induced αs1-casein gene expression in rabbit primary mammary cells: Role of STAT5, C/EBP, and chromatin structure. J. Cell. Biochem. 95, 313–327 (2005).

27. Maharana, S. et al. Chromosome intermingling—the physical basis of chromosome organization in differentiated cells. Nucleic Acids Res. 44, 5148–5160 (2016).

28. Wang, Y., Nagarajan, M., Uhler, C. & Shivashankar, G. V. Orientation and repositioning of chromosomes correlate with cell geometry-dependent gene expression. Mol. Biol. Cell 28, 1997–2009 (2017).

29. Ghosh, S. et al. Dedifferentiation alters chondrocyte nuclear mechanics during in vitro culture and expansion. Biophys. J. (2021) doi:10.1016/J.BPJ.2021.11.018.

30. Maki, K. et al. Hydrostatic pressure prevents chondrocyte differentiation through heterochromatin remodeling. J. Cell Sci. 134, (2021).

31. Murphy, M. K., Huey, D. J., Hu, J. C. & Athanasiou, K. A. TGF-β1, GDF-5, and BMP-2 Stimulation Induces Chondrogenesis in Expanded Human Articular Chondrocytes and Marrow-Derived Stromal Cells. Stem Cells 33, 762–773 (2015).

32. Schneider, M. C., Chu, S., Randolph, M. A. & Bryant, S. J. An in vitro and in vivo comparison of cartilage growth in chondrocyte-laden matrix metalloproteinase-sensitive poly(ethylene glycol) hydrogels with localized transforming growth factor β3. Acta Biomater. 93, 97–110 (2019).

33. Mikkelsen, T. S. et al. Dissecting direct reprogramming through integrative genomic analysis. Nature 454, 49–55 (2008).

34. Stratton, M. S. & McKinsey, T. A. Epigenetic regulation of cardiac fibrosis. J. Mol. Cell. Cardiol. 92, 206–13 (2016).

35. Heerboth, S. et al. Use of epigenetic drugs in disease: an overview. Genet. Epigenet. 6, 9–19 (2014).

36. Lui, J. C. et al. EZH1 and EZH2 promote skeletal growth by repressing inhibitors of chondrocyte proliferation and hypertrophy. Nat. Commun. 7, 13685 (2016).

37. Yamamoto, H., Schoonjans, K. & Auwerx, J. Sirtuin Functions in Health and Disease. Mol. Endocrinol. 21, 1745–1755 (2007).

